# Unique features of the arterial Blood-Brain Barrier

**DOI:** 10.1101/2022.10.30.514417

**Authors:** Batia Bell, Shira Anzi, Esther Sasson, Ayal Ben-Zvi

**Author notes:** **Address correspondence to:** Ayal Ben-Zvi, PhD, Department of Developmental Biology and Cancer Research, The Institute for Medical Research Israel-Canada, Faculty of Medicine, Hebrew University of Jerusalem, Jerusalem 91120, Israel.

## Abstract

CNS vasculature differs from vascular networks of peripheral organs by its ability to tightly control selective material exchange across capillary barriers. Capillary permeability is mostly defined by unique cellular components of the endothelium. While capillaries are extensively investigated, the barrier properties of larger vessels are understudied. Here, we investigate barrier properties of CNS arterial walls. Using tracer challenges and various imaging modalities, we discovered that at the mouse cortex, the arterial barrier does not reside at the classical level of the endothelium. The arterial wall’s unique permeability acts bi-directionally; CSF substances travel along the glymphatic path and can penetrate from the peri-vascular space through arteriolar walls towards the lumen. We found that caveolae vesicles in arteriole endothelial and smooth-muscle cells are functional transcytosis machinery components, and that a similar mechanism is evident in the human brain. Our discoveries highlight vascular heterogeneity investigations as a potent approach to uncover new barrier mechanisms.

## Main

Organs receive nutrients and clear metabolic waste through the blood. Vascular networks are largely assembled in a stereotypic manner in which the smallest diameter vessels (capillaries) are constructed by an inner lining of very thin endothelial cells (one or two cells in each tube section) surrounded by pericytes. The thinnest vessel wall and the very slow blood flow facilitate metabolic material exchange between the blood and the tissue. In general, arteries and veins facilitate blood supply and drainage, but each also have additional functions. Larger diameter vessels are constructed by an inner lining of endothelial cells surrounded by layers of smooth muscle (and other cell types). Arteries and their bifurcations (arterioles) function by diverging blood flow to different parts of the network by virtue of smooth muscle driven vessel constriction and dilation.

In the central nervous system (CNS), cellular processes of astrocytes (termed end-feet) construct a third layer covering each vessel. CNS vasculature also differs from peripheral organs’ vascular networks in its ability to tightly control selective metabolic material exchange across capillary walls. This and additional features, such as preventing immune cells entry into the CNS, and active clearance of toxins from the tissue, are hallmarks of the blood-brain barrier (BBB)^1,2^.

The degree of capillary permeability (i.e. the ease by which blood-borne materials extravasate into the tissue) is largely set by unique cellular components of the endothelium. As such, the basic components of the BBB are continuous endothelial cells lacking fenestrations, negatively suppressing vesicular transport and constructing highly impermeable tight junctions^1,3^. Based on tracer studies, endothelial BBB properties are assumed to exist at every level of the vascular network^1^. Nonetheless, endothelial barrier properties are mainly investigated at the capillary level, while barrier properties of larger vessels are understudied.

Emerging evidence of endothelial heterogeneity across CNS vessel types^4^ prompted us to investigate barrier properties of CNS arterial walls. We used tracer challenges and different imaging modalities and discovered that at the mouse cortical arterial wall, the BBB does not reside at the level of the endothelium. Despite relatively fast blood flow, we found non-selective penetrance of various substances from the blood, crossing both the endothelial and smooth muscle layers, but not the astrocyte end-feet layer. Arterial wall unique permeability acts bi-directionally; substances originating from the CSF can penetrate from the perivascular space, crossing the smooth muscle layer and the endothelial layer towards the vessel lumen. We show that caveolae vesicles in arteriole endothelial and smooth muscle cells are functional transcytosis machinery components, and that a similar mechanism is also evident in endothelial and smooth muscle cells of human tissue.

## Results

### Atypical cellular barrier properties at the CNS arteriole-wall

Tracers of different sizes and molecular compositions are commonly used to test vascular permeability. We introduced tracers into the blood stream of wild-type adult mice for relatively short permeability challenges (10 min), followed by immunostaining for Neuro-vascular unit (NVU) components in cortical sections (endothelium, smooth muscle, astrocyte end-feet, and basement membranes). Cortical arterioles usually have a circumferential organization of a single endothelial layer, a smooth muscle layer and astrocyte end-feet, each separated by a basement membrane (Figure 1a, illustration). We focused mostly on arterioles (of 5-10 μm diameter), but the following observations are also relevant to larger diameter cortical-penetrating arteries. With confocal microscopy imaging we found differing extents of permeability along the arteriole wall for several tracers. Albumin (70 kDa) was mostly co-localized with the endothelium (Figure 1b). Dextran (10 kDa) signals co-localized with the endothelium and reached the smooth muscle layer, and in some cases reached beyond, up to the astrocyte end-feet (Figure 1c). Sulfo-biotin (443 Da) signals localized past the smooth muscle layer, but were confined by the outer basement membrane (Figure 1d).

**Figure 1.**
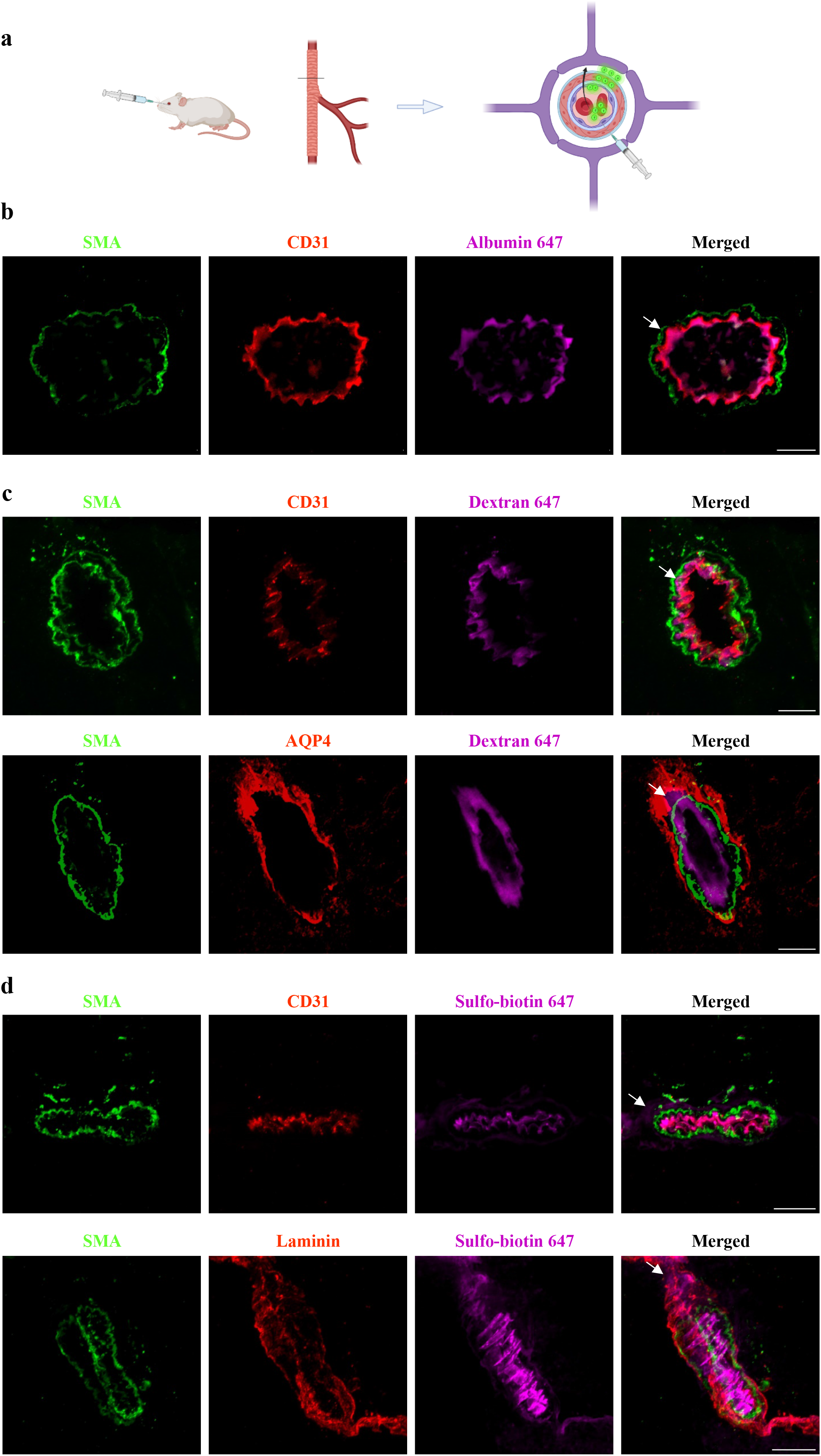
Variable degree of permeability to different tracers across the CNS arteriole-wall uncovers atypical cellular barrier properties. Confocal-microscopy of arteriole-wall permeability, with immunostaining for NVU components: endothelium (anti-CD31), smooth-muscle (anti-SMA), astrocyte end-feet (anti-AQP4), and basement membranes (anti-pan Laminin) of wild-type adult mouse cortical sections. **a**, Schematic illustration of the experimental design: tracers of different size and molecular compositions are introduced into the blood stream and circulated for 10 min. Arteriole cross sections are used to locate fluorescent tracer signals (green circles) in the vessel wall from luminal side (blood) towards the tissue (arrow showing the path crossing the endothelial cell layer-purple, smooth-muscle layer-red each surrounded by basement membranes and all surrounded by the astrocyte end-feet) Created with BioRender.com. **b**, Tracer challenges with Alexa-647 conjugated albumin (70 kDa) demonstrating that albumin is mostly co-localized with the endothelium (arrow). **c**, Challenges with Alexa-647 conjugated dextran (10 kDa) demonstrates tracer signals co-localized with the endothelium and reaching the smooth-muscle layer (upper panel, arrow), and beyond reaching the astrocyte end-feet (lower panel, arrow). **d**, Challenges with sulfo-biotin (443 Da), stained with Alexa-647 conjugated streptavidin demonstrates tracer signals passed the smooth-muscle layer (upper panel, arrow), but confined by the basement membranes (lower panel, arrow). Images are representative of n=27 arterioles profiles of n=9 mice (3 for each tracer), Scale Bars 10um.

The juxtaposed organization of thin cell profiles in vessel cross-sections makes it difficult to determine precise localization of tracer molecules with limited imaging resolution. We therefore used super-resolution imaging (dSTORM). Precise nano-scale localization of single tracer molecules confirmed localization past both the endothelium and the smooth muscle markers of all three tracers (Figure 2, S1). Permeability quantification showed a substantial percentage of tracer molecules beyond the smooth muscle marker limits, with no significant difference between the three tracers tested (Figure 2c). With appropriate imaging resolution, at the most abluminal location we could determine that tracers were arranged in a confined manner and did not freely disperse further into the tissue.

**Figure 2.**
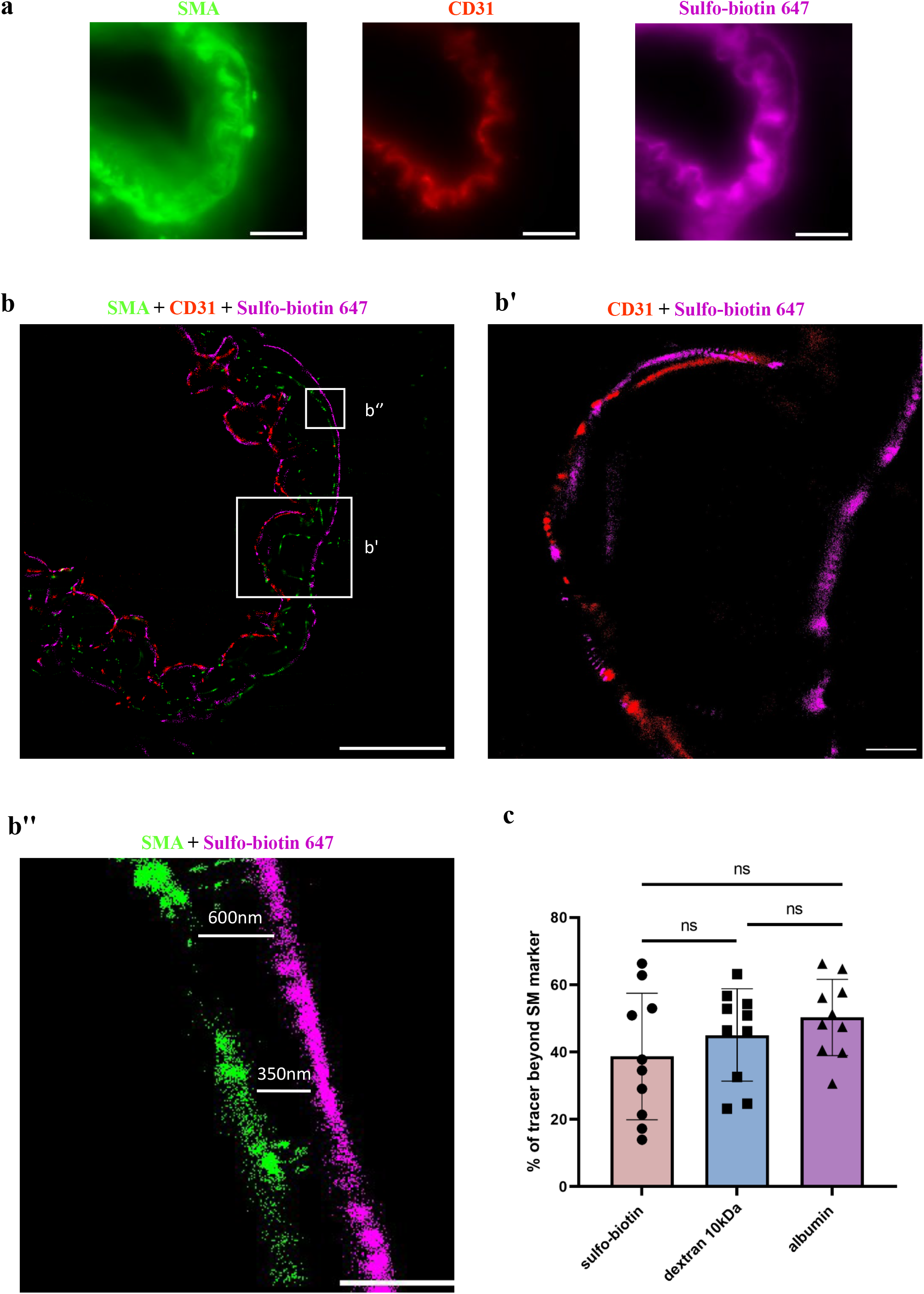
STORM imaging demonstrates super-resolution tracer permeability and validates the unique barrier properties of the CNS arteriole wall. Precise nano-scale localization of the sulfo-biotin tracer (443 Da), stained with Alexa-647 conjugated streptavidin. **a**, Relatively low resolution TIRF imaging of endothelium (anti-CD31), smooth-muscle (anti-SMA) in wild-type adult mouse brain sections, does not allow precise localization of tracer along the vascular wall due to diffraction limitation. Scale bar 10 μm. **b**, STORM images of the same arteriole (as in **a**) shows that tracer signals are found passed both the endothelium and the smooth-muscle markers. Scale bar 10 μm. **b’-b’’**, Inset magnifications showing distances between the tracer and the cell markers (CD31 (**b’**) and of SMA (**b’’**), scale bars b’ 2 μm, b’’ 1 μm). **c**, Quantification of tracer signals beyond the smooth muscle marker (sulfo-biotin tracer images appear here, dextran and albumin tracers images appear in Figure S1). There are no significant differences in permeability of these three tracers of different size and molecular compositions (Kruskal–Wallis H test). Images are representative of n=18 arteriole profiles of n=4 mice.

We conclude that unlike brain capillaries (Figure S2), the BBB in arterial walls does not reside at the level of the endothelium -in mouse CNS arteries various substances can penetrate from the blood, crossing the endothelial layer and the smooth muscle layer, but not the astrocyte end-feet layer.

### Caveolae vesicles in arteriole endothelial and smooth-muscle cells are functional transcytosis machinery components

Recently, Chow *et al*. found that caveolae-vesicles in CNS arterioles mediate neurovascular coupling (arteriole constriction/dilation in response to neuronal activity)^5^. Vesicles were found in both the endothelium and the smooth muscle cells. Elegant experiments of genetic perturbations that eliminated these components revealed that only the endothelial vesicles are necessary for neurovascular coupling^5^. Since suppression of endothelial caveolae-vesicles formation is a hallmark of capillary BBB endothelium^3,6-8^ (preventing non-specific transcytosis across the barrier), we wondered if arteriole caveolae vesicles could be transcytotic components mediating the unique permeability pattern that we described. To answer this question, we used electron microscopy to image permeability challenges of two tracers: sulfo-biotin (443 Da), and horseradish peroxidase (HRP, ∼40 kDa). Both tracers could be found in basement membranes between all three cell layers, confirming permissive permeability (Figure 3, arrows). Abundant vesicular structures were observed in both the endothelium and the smooth muscle cells, and tracers could be found filling these vesicles (Figure 3, arrowheads). Tracers were found in luminal and abluminal membranes pits, and in the cytoplasm of both cell types. We concluded that intense vesicular activity in these cells reflects potential cargo trafficking, and therefore supports functional transcytosis machinery that underlie the unique arteriole permeability pattern.

**Figure 3.**
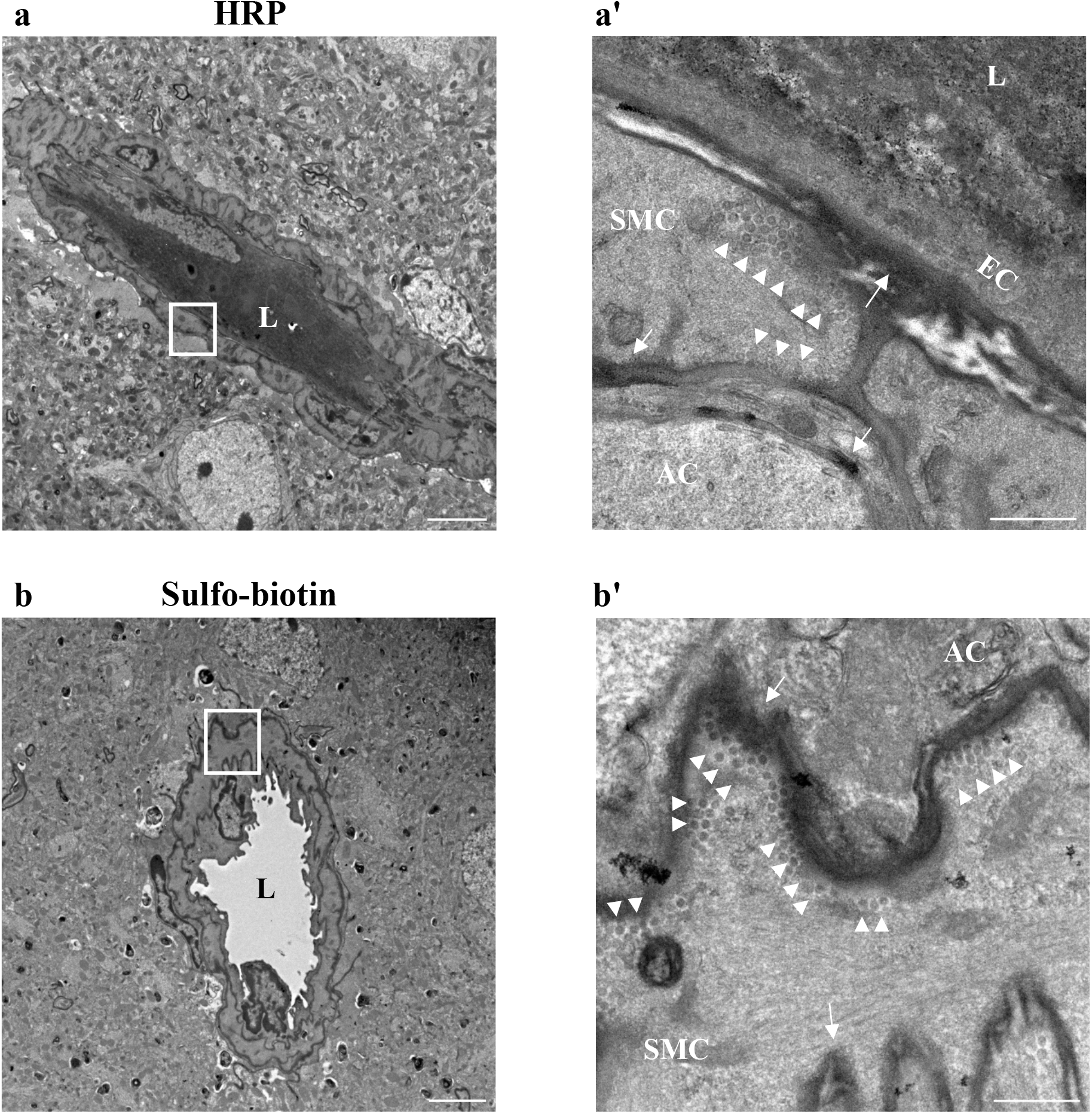
Caveolae vesicles in arteriole endothelial and smooth muscle cells are functional transcytosis machinery components. HRP and sulfo-biotin tracers, imaged by TEM, demonstrate cargo trafficking in CNS arteriole cells. **a**, Representative TEM image of a cortical arteriole (wild-type adult mice) following HRP tracer challenge. HRP signal is found in vessel lumen (L) as well as in basement membranes; in between the endothelial layer and smooth muscle layer (arrows), and between the smooth-muscle layer and astrocyte end-feet (arrows). Scale bar 5 μm. Inset (**a’**) ample caveolae vesicles, some of which showing HRP signals at luminal and abluminal membranes of a smooth muscle cell (arrowheads). Scale bar 500 nm. **b**, Representative TEM image of sulfo-biotin tracer challenge following staining with HRP conjugated streptavidin. Scale bar 5 μm. Inset (**b’**) high magnification image showing abundant tracer-field vesicles, adjacent to the abluminal membrane of a smooth-muscle cell (arrowheads). Tracer signal is found also in basement membranes (arrows). SMC-Smooth-muscle cell, EC- Endothelial cell, AC-Astrocyte, L-lumen. Scale bar 500 nm (n = 6 mice, 12 arterioles).

### Arterial wall permeability is bi-directional

Our electron microscopy study supports transcytosis-based permeability from the blood into the arteriole wall. Nonetheless, it could not determine the directionality of cargo movement in other axes. To address this issue, we used confocal and super-resolution imaging, this time introducing tracers into the CSF. A ‘glymphatic’ path describes movement of CSF from the sub-arachnid space, along the perivascular space created between the astrocyte end-feet in the glia limitans (continuous with these covering the parenchymal side of the Pia, at the Virchow-Robin spaces of penetrating arteries) and the smooth muscle layer^9^ (Figure 4a, illustration). Adopting the methodology used to study this path^9^, we injected dextran (10 kDa) tracer into the cisterna magna. First, we confirmed that tracers reached the perivascular spaces around arterioles (Figure 4b-c, confocal microscopy), in which we could find approximate co-localization of tracers with both endothelium and smooth muscle signals. With super-resolution imaging (dSTORM), we could show precise nano-scale localization of single tracer molecules past the smooth muscle marker (Figure 5a, arrows) and at the luminal side of the endothelium (arrowhead, the blood side). With this approach, we could see that tracers originating in the CSF compartment end up in the blood compartment across the arteriole wall. Based on tracer signal distribution, we identified tracer clusters that match the dimensions of transcytosis vesicles (Figure 5b’). We also found vesicle-shaped structures connected to the basement membrane that resemble the ultrastructure of flask-shape membrane pits (Figure 5b’). Therefore, we concluded that arteriole permeability is bi-directional, probably mediated also by trancytosis.

**Figure 4.**
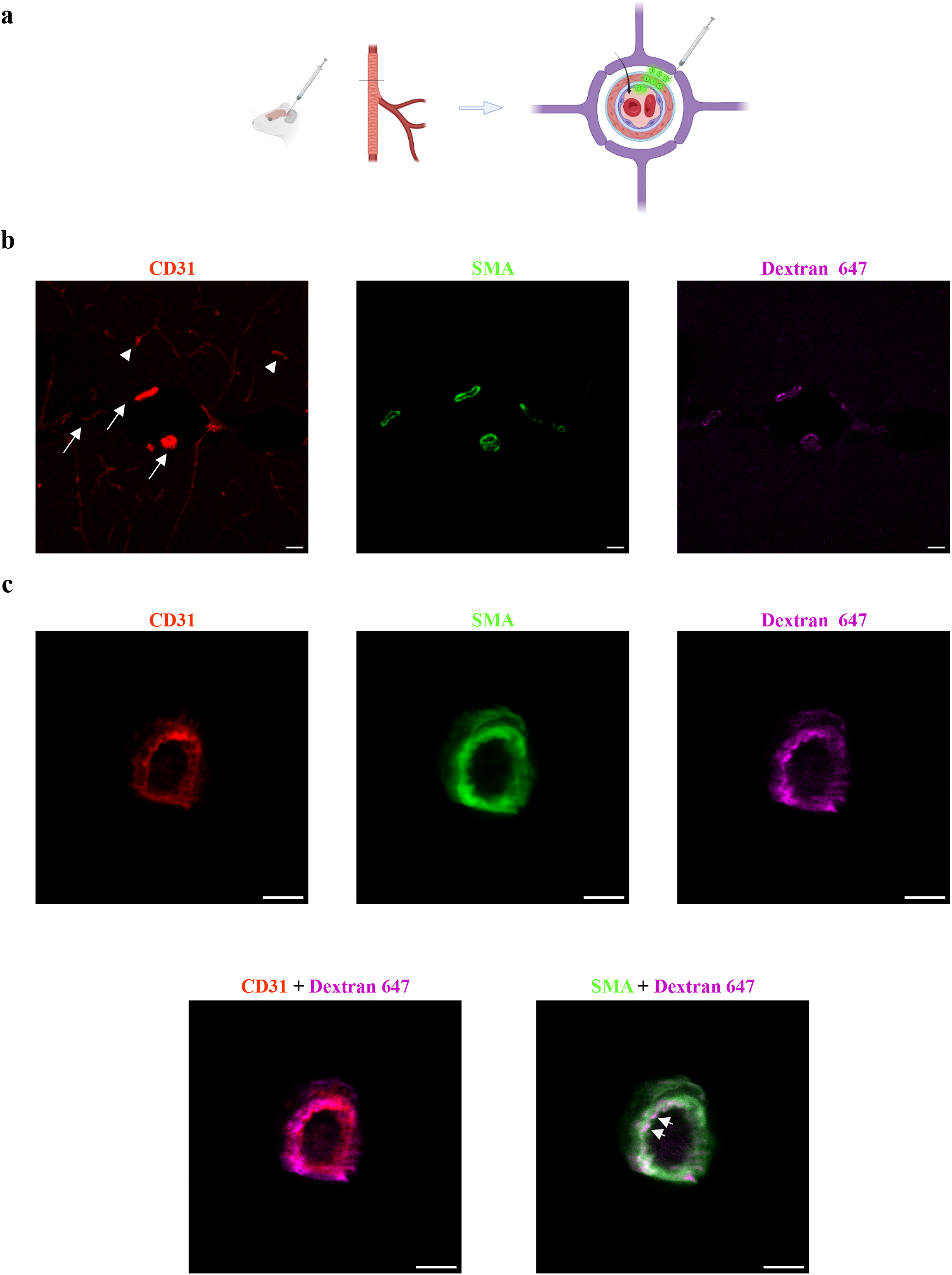
Tracer introduced into the CSF penetrates into the perivascular space of arteriolar wall. For testing the potential of clearance mechanisms, we adopted the methodology used to study the ‘Glymphatic’ path, and injected Dextran (10 kDa) into the cisterna magna. **a**, Schematic illustration of the experimental design: very small volumes of tracers are introduced into the cisterna magna and circulate in the CSF for 30 min. Cortical cross sections are used to locate fluorescent tracer signals (green circles). **b**, Low magnification confocal microscopy of cortical sections demonstrates expected tracer signals around arteries/arterioles (arrows, CD31+SMA double positive vessels), but not around capillaries (arrowheads, CD31 single positive vessels). Scale bar 5 μm **c**, Confocal microscopy imaging of an arteriole cross section demonstrates approximate co-localization of the tracer with both endothelium and smooth-muscle signals (arrows). Scale bar 10 μm (n = 3 mice, 12 arterioles).

**Figure 5.**
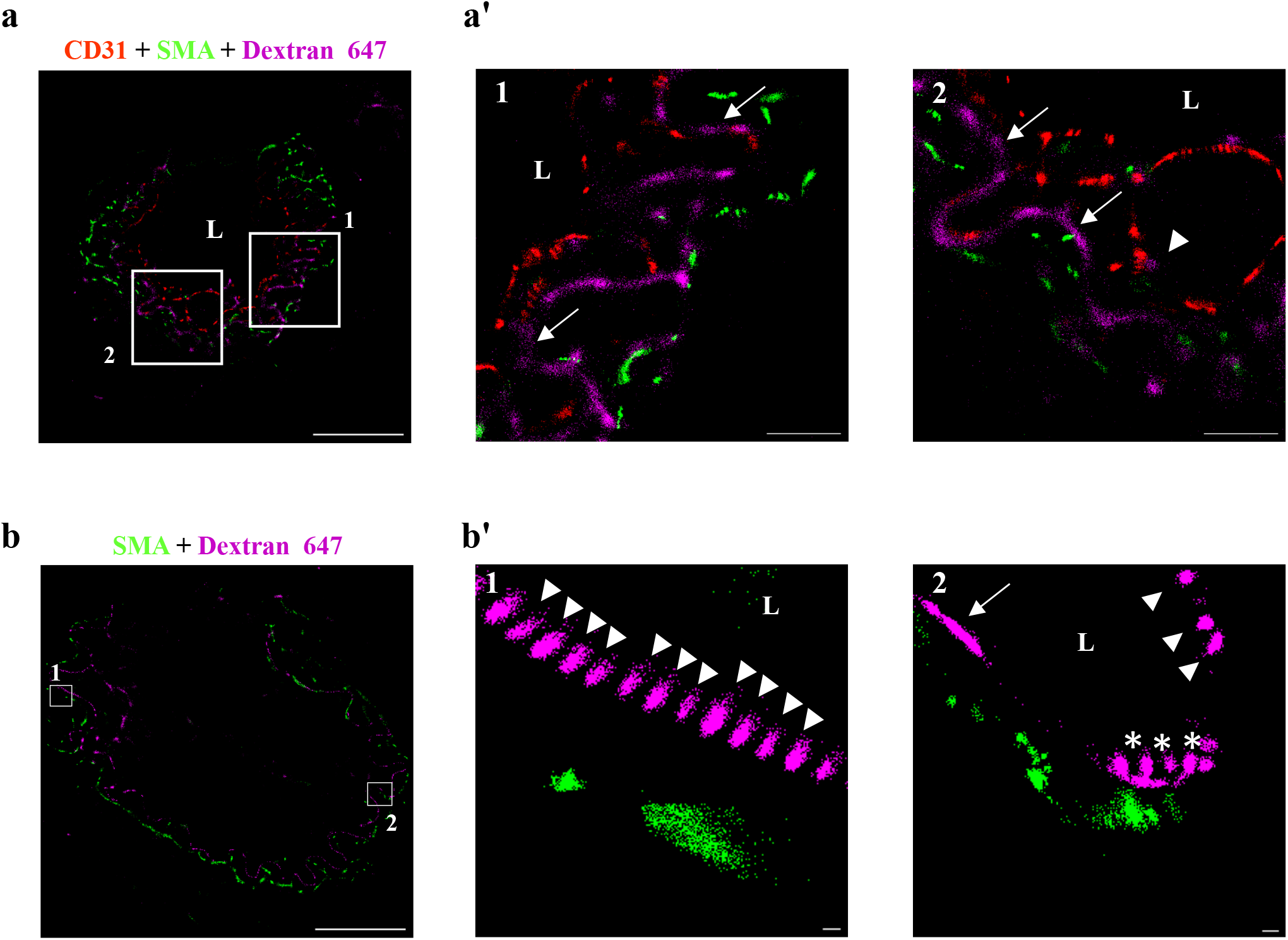
Arterial wall permeability is bi-directional. Tracer introduced into the CSF travel along the Glymphatic path and can penetrate from the perivascular space across arteriolar walls towards the luminal direction. **a-b**, CSF tracer challenges with Alexa-647 conjugated dextran (10 kDa, cisterna magna injections) and immunostaining for SMA and CD31 of wild-type adult brain sections (L-vessel lumen). Scale bars 10 μm **a**, Precise nano-scale localization with STORM imaging shows tracer signals located along the CSF-blood trajectory. Inset (**a’** scale bars 2 μm) magnifications demonstrating tracer signals between smooth-muscle and endothelium markers (arrows), and at the endothelium luminal side (arrowhead). **b**, Based on tracer signal distribution, especially at high magnification (inset **b’** scale bars 100 nm), we identify tracer clusters that fit the dimensions of transcytosis vesicles. These vesicle-like structures are located in smooth muscle cells (arrowheads, in 1 and 2). Elongated distribution might represent tracer filled basement membrane between the smooth muscle and the endothelial cell (arrow, in 2). Vesicle-shaped structures that are connected to the basement membrane resemble the ultrastructure of flask shape membrane pits (astrix); n = 3 mice, 12 arterioles.

### Evidence of transcytosis in human brain arterioles

The H01 dataset is a 1.4 petabyte rendering of a small sample of human brain tissue (one cubic millimeter), released by a collaboration between the Lichtman Laboratory at Harvard University and Google^10^. A rapidly preserved human surgical sample from the temporal lobe of the cerebral cortex was imaged at nanoscale-resolution by serial section electron microscopy, reconstructed and annotated by automated computational techniques^10^. We used this dataset to search for transcytosis components in human cortical vasculature. Our analysis shows that properties of the human vasculature correspond with the mouse data (figure 3 and previous published data^5^). In the one cubic millimeter of human temporal lobe, we found 5 arterioles. Examining these vessels along the serial section electron microscopy reconstruction, we found extensive vesicular activity in both endothelium and smooth muscle cells in patterns that resembles the mouse data. Quantifying vesicular density showed significant higher vesicular density in arterial endothelium than in capillary endothelium (Figure 6).

**Figure 6.**
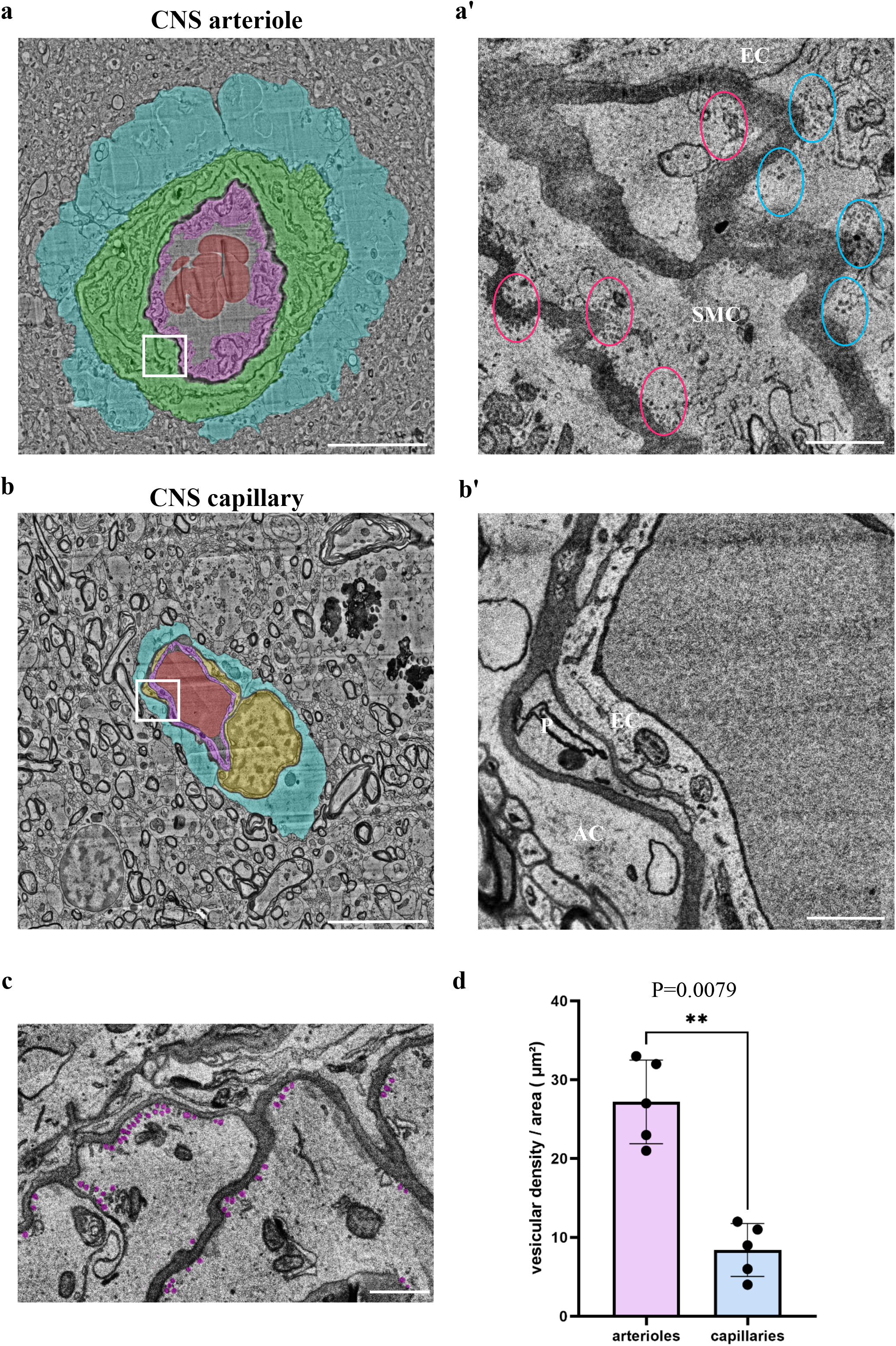
Abundant vesicles in arteriole cells of human CNS might indicate transcytotic activity. Caveolae vesicles in arteriole endothelial and smooth muscle cells are evident in 3D EM reconstruction of human cortical tissue^10^. **a-b**, Transmission electron microscopy images of a CNS arteriole and capillary. Pseudo-colors highlight cell types: smooth-muscle (green), endothelium (purple), pericyte (yellow), astrocyte end-foot (blue), red blood cell (red), Scale bar 10 μm. **a’-b’**, Magnification of the boxed area in the left panel. **a’**, Abundant vesicles in aEC’s (blue ovals) and aSMCs (red ovals). **c**, Example of multiple vesicles (pseudo labeled in magenta) in aSMCs. **d**, Quantification showing significant higher vesicular density in aECs than in cECs (n=5 arterioles and 5 capillaries). SMC-Smooth-muscle cell, EC-Endothelial cell, AC-Astrocyte, P-Pericyte, L-lumen. Scale bar 1 μm. Data are mean ± s.e.m. *p < 0.05 (two tailed Mann–Whitney U-test).

## Discussion

Our study reveals distinct cellular barrier properties in different segments of the CNS vascular tree. Unlike brain capillaries, the BBB in arterial walls does not reside at the level of the endothelium and various substances can penetrate from the blood, crossing the endothelium and the smooth muscles, while being restricted from crossing the astrocyte end-feet layer. It is possible that unique barrier properties of arterioles were overlooked simply because barrier properties are mostly studied at the vascular site where physiological material exchange occurs (capillaries). In this context, it is surprising that fast blood flow allows this permeability even in relatively short-term tracer challenges. Based on our current study, hyper-permeability is at least in part mediated by vesicular transport, but we cannot exclude involvement of other pathways, and future studies should also examine the possibility of unique structure and function of arteriole tight junctions.

We show here that the recently discovered caveolae vesicles in arteriole endothelial and smooth-muscle cells^5^ are functional transcytosis machinery components. Vesicular activity is dependent on the expression of Cav1 (a structural vesicle component) in both cell types, but only the endothelial activity has been linked to neurovascular coupling^5^. Since we show that these vesicles carry cargo from the blood, future studies might focus on testing a possible involvement of specific cargoes in modulation of neurovascular coupling. At the molecular level, it is known that in capillary endothelium, the expression of Mfsd2A (a BBB specific lipid transporter) negatively regulates transcytosis^3,6,7^. Interestingly Mfsd2A mRNA is abundant also in arteriole endothelium^4^, but is absent at the protein level^5^. Forced transgenic Mfsd2a expression suppresses vesicular activity and abrogates neurovascular coupling^5^.

We were fascinated by the observation that the arterial wall’s unique permeability properties are bi-directional - substances originating in the CSF can penetrate from the perivascular space, crossing the smooth muscle layer and the endothelial layer towards the blood side. We found preliminary evidence that this movement is at least in part mediated by reverse transcytosis (Figure 5). We suggest two hypotheses for the physiological function of this phenomenon: endothelial vesicular activity is obligatory for neurovascular coupling, but as a by-product it affects barrier selectivity in arterioles. Reverse transcytosis of smooth muscle cells might have evolved in order to compensate for this non-selective leakage. Alternatively, regardless of neurovascular coupling, active reverse transcytosis mechanisms might function in the clearance of materials from the CSF into the blood.

This unique arterial BBB structure can also be found in the human brain - arteriolar specific caveolae vesicles are evident in endothelial and smooth muscle cells of human tissue. In this analysis we observed vesicles also in other large vessels, presumably veins. Permeability and function of these structures should be explored in future studies, especially in light of the importance of veins and postcapillary venules in mediating immune cell extravasation^11^. Moreover, a recent study found that transferrin receptor-targeted liposome nano-particles for the use of drug delivery preferably exploit transport at postcapillary venules^12^.

Finally, astrocyte end-feet have been suspected for many years to be anatomical counterparts of the BBB. The discovery of an endothelial barrier diminished the focus of studying atrocyte end-feet barrier functions. From an evolutionary perspective, invertebrate glia cells function as CNS barriers. Even ancestral vertebrates had glial barriers (elasmobranch fish like sharks, skates, and rays display this feature^13^). Along with the appearance of vascular systems, barrier functions shifted to the endothelium, and an endothelial barrier became dominant in all vertebrates^13^. Endothelial barrier evolution does not necessitate the degeneration of the glial barrier. Technically, it is hard to devise *in vivo* assays to test end-feet potential barrier functions, since blood-borne tracers are sequestered already at the capillary endothelial barrier. We show that arterioles’ astrocyte end-feet layer is the last point of tracer penetrance from the blood. At this stage we have no direct evidence for cellular or molecular barrier properties of astrocytes and we are not aware of known molecular differences between arteriole and capillary associated astrocytes. Another incentive to look for such functions is that anatomically, arteriole astrocyte end-feet are glia-limitans, continuous with astrocyte end-feet covering the parenchymal side of the Pia, which possesses barrier properties. Uncovering molecular components of this putative barrier might provide new targets for drug delivery through the arterial path. We shed light on the potential of substance exchange in this previously unappreciated vascular site. Altogether we expect that the concept of a single unifying BBB will be revised, and discoveries of heterogeneity in barrier mechanisms will open new avenues for drug delivery and for better understanding the physiology of CNS vasculature.

## Methods

### Mice

All mice were maintained in the animal facility of the Hebrew University under specific pathogen-free conditions. All animals were treated according to institutional guidelines approved by the Institutional Animal Care and Use Committee (IACUC) at Hebrew University (protocol MD-21-16361-5). 8-9 week old male and female C57BL/6JOlaHsd and ICR mice were purchased from Envigo (Rehovot, Israel).

### Tissue preparation

After dissection, brains were fixed in 4% paraformaldehyde (PFA, Sigma Aldrich) at 4 °C overnight, cryopreserved in 30% sucrose and frozen in TissueTek OCT (Sakura). Frozen brains were cut to either 10 μm slices for immunofluorescence staining or 4-6 μm for STORM (CM1950, Leica) to produce coronal brain sections.

### Immunofluorescence

10 μm thick cryo-sections were washed with phosphate buffered saline (PBS) for 5 min at room temperature (RT) and then incubated for 3 h at room temperature (RT) with blocking solution (10% goat serum (GS), 10% horse serum (NHS), 0.05% triton X-100 in PBS). Slides were incubated with primary antibodies (diluted in 2.5% GS, 2.5% NHS, 0.05% triton X-100 in PBS) at 4 °C overnight (see antibodies table for details; Table 1). Slides were then washed with PBS, incubated with secondary antibodies for 1 h at RT and washed again. Samples were mounted with freshly made imaging buffer for dSTORM (describe in the dSTORM imaging section) and epifluorescence microscopy, or mounted in Fluoromount G (EMS) for confocal microscopy.

**Table 1:**
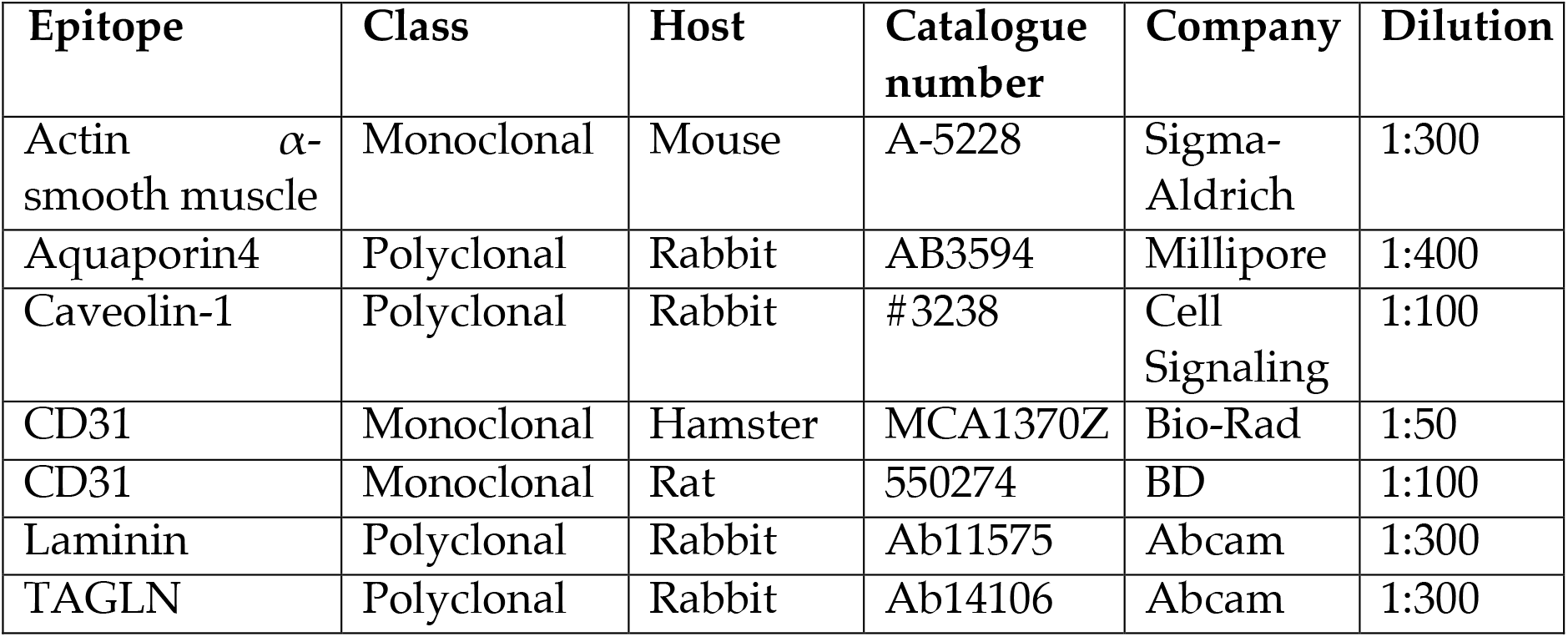
Details of antibodies used

| Epitope | Class | Host | Catalogue number | Company | Dilution |
| --- | --- | --- | --- | --- | --- |
| Actin α-smooth muscle | Monoclonal | Mouse | A-5228 | Sigma-Aldrich | 1:300 |
| Aquaporin4 | Polyclonal | Rabbit | AB3594 | Millipore | 1:400 |
| Caveolin-1 | Polyclonal | Rabbit | #3238 | Cell Signaling | 1:100 |
| CD31 | Monoclonal | Hamster | MCA1370Z | Bio-Rad | 1:50 |
| CD31 | Monoclonal | Rat | 550274 | BD | 1:100 |
| Laminin | Polyclonal | Rabbit | Ab11575 | Abcam | 1:300 |
| TAGLN | Polyclonal | Rabbit | Ab14106 | Abcam | 1:300 |

**Table 2:** Details of fluorophores used

| Fluorophore | Isotype | Catalogue number | Company | Dilution |
| --- | --- | --- | --- | --- |
| Alexa fluor488 | Anti-mouse IgG | 715-545-151 | Jackson | 1:800 |
| Alexa fluor568 | Anti-rabbit IgG | A11011 | Life Technologies | 1:1000 |
| Alexa fluor647 | Anti-rat IgG | 712-605-153 | Jackson | 1:1200 |
| Atto488 | Anti-mouse IgG | 62197 | Sigma-Aldrich | 1:800 |
| Atto488 | Anti-rabbit IgG | 18772 | Sigma-Aldrich | 1:800 |
| CF568 | Anti-mouse IgG | 20105 | Biotum | 1:800 |
| CF568 | Anti-rabbit IgG | 20098 | Biotum | 1:800 |
| Streptavidin Alexa fluor647 | Biotin | S32357 | Molecular Probes | 1:1000 |
| Streptavidin HRP | Biotin | S911 | Invitrogen | 1:1000 |

### BBB permeability assay

In all experiments, a leakage incident was defined when a tracer was localized outside the endothelial or smooth muscle cell area (indicated by marker or ultrastructure). Relative leakage index was calculated as tracer signal density (signals/area) in an arbitrary fixed area and distance from the luminal side of the relevant marker.

### Dextran and albumin

Deeply anesthetized (8.5 mg/ml ketamine, 1.5 mg/ml xylazine, in 100 μl saline) 8-9 week old male and female C57BL/6JOlaHsd mice were injected retro-orbitally with 20 μl of Alexa Fluor647 anionic fixable conjugated Dextran (D22914, Molecular Probes, 4mg/ml) or 120 μl of Alexa Fluor647 conjugated BSA (A34785, Thermo-Fisher, 5mg/ml). The tracer was allowed to circulate for 10 min. Brains were dissected and handled as described under tissue preparation section.

### NHS-biotin

Deeply anesthetized 8-9 week old male and female C57BL/6JOlaHsd mice were perfused for 5 min with sulfo-NHS-biotin (4 mg/20 g mouse body weight, Thermo Scientific, cat no. 21217, dissolved in 20 ml PBS). Brains were dissected and handled as described under tissue preparation section.

### HRP

Deeply anesthetized 8-9 week old male and female ICR mice were injected retro-orbitally with 0.4 ml of HRP (Sigma Aldrich, HRP type II, 10 mg/20 g mouse body weight, dissolved in PBS). After 30 min of HRP circulation, brains were dissected and fixed as described in the TEM section.

### Intracisterna magna (i.c.m.) injection

Mice were anesthetized with ketamine/medetomidine i.p. and injected i.c.m. into the cisterna magna with 2 μl of Dextran, Alexa Fluor647 anionic, fixable (D22914, Molecular Probes, 4 mg/ml). Mice were then left on a heating pad for 30 min before brains were dissected and handled as described in the tissue preparation section.

### Transmission electron microscopy (TEM)

Freshly dissected tissue samples were fixed with Karnovsky’s fixative (2% PFA, 2.5% glutaraldehyde in 0.1 M cacodylate buffer, pH = 7.4) for 4 h at RT, followed by a 1:2 dilution of Karnovsky’s fixative in 0.1 M cacodylate buffer overnight at 4 °C. Sections of 60–80 nm were cut on Leica vibrating blade microtome (VT1000S), and developed with DAB. For 3,3′-Diaminobenzidine (DAB) staining, sections were incubated in 0.05 M Tris-HCl buffer (pH 7.6) containing 5 mg 3-3′ diaminobenzidine (Thermo Scientific, TA-060-HDX) per 10 mL buffer and a final concentration of 0.01% hydrogen peroxide for 15 min at RT. The DAB reaction was stopped with PBS wash and quenched by post fixation in 2% osmium tetroxide sodium cacodylate buffer (OsO4). Samples were post-fixed in 1%OsO4 in 0.1 M cacodylate buffer for 2 h, dehydrated in a graded series of alcohols, and embedded in epoxy resin. Sections of 60–80 nm were cut on an ultramicrotome (Ultracut, Reichert-Jung) contrasted with uranyl acetate and lead citrate, and examined with a Jeol (JEM-1400 PLUS, Japan) electron microscope.

### Fluorescence microscopy

Immunofluorescence images were captured using the following confocal microscopes: Nikon Eclipse Ni, objective ×20 and ×40 with a Nikon C2 camera, and Nis-Elements software, or with Nikon TE-2000, objective ×20, ×40 and ×60 with EZ-C1 software.

### dSTORM imaging

We used a dSTORM system, which allows imaging at approximately 20 nm resolution by using photo-switchable fluorophores (all dSTORM imaging was done on TIRF mode). 4-6 μm brain slices were mounted on poly-D-lysine coated coverslips (no. 1.5 H, Marienfeld-superior, Lauda-Königshofen, Germany). dSTORM imaging was performed in a freshly prepared imaging buffer containing 50 mM Tris (pH 8.0), 10 mM NaCl and 10% (w/v) glucose with an oxygen-scavenging GLOX solution (0.5 mg/ml glucose oxidase (Sigma-Aldrich)), 40 μg/ml catalase (Sigma-Aldrich), 10 mM cysteamine MEA (Sigma-Aldrich), and 1% β mercaptoethanol (Barna et al., 2016; Dempsey et al., 2011; Zhang et al., 2016)^14-16^.

A Nikon Ti-E inverted microscope was used. The N-STORM Nikon system was built on TIRF illumination using a 1.49 NA X100 oil immersion objective and an ANDOR DU-897 camera. 488, 568 and 647 nm laser lines were used for activation with cycle repeat of

∼8000 cycles for each channel. Nikon NIS Element software was used for acquisition and analysis; analysis was also performed by ThunderSTORM (NIH ImageJ [Ovesný et al., 2014]^17^).

### dSTORM quantifications

The dSTORM approach we used is based on labeling the target protein with a primary antibody and then using a secondary antibody conjugated to a fluorophore. Thus, resolved signals represent a location that is approximately 40 nm from the actual epitope (assuming the approximation of the two antibodies’ length in a linear conformation). The number of signals represents an amplification of the actual target numbers. Amplification corresponds to the primary antibody in the case of a polyclonal antibody (assuming binding to several epitopes in the same protein, which could be reduced by the use of monoclonal antibodies). Amplification also corresponds to several secondary antibodies binding to a single primary antibody and to several fluorophores attached to a single secondary antibody. Nevertheless, resolution of approximately 20 nm allows us to separate signals and to use these as proxies to the abundance of target molecules, which can reliably be used to compare different states^18,19^.

Single molecule localization microscopy (SMLM) results in point patterns having specific coordinates of individual detected molecules. These coordinates are typically summarized in a ‘molecular list’ (provided by ThunderSTORM analysis (NIH ImageJ) [Ovesný et al., 2014]^17^).

### Statistical analysis

All comparisons were performed by two-tailed Mann–Whitney U-tests, (as indicated in the figure legends), for multiple comparisons Kruskal–Wallis H test was used, p < 0.05 was considered significant. (GraphPad Prism 8.0.1 [244] for Windows, GraphPad Software, San diego, California, USA).

**Illustrations were created with BioRender.com**

## Acknowledgments

We would like to thank; the Ben-Zvi group for scientific inputs, Dr. Evellina Sjostedt and Dr. Yaron Meirovitch for scientific inputs and guidance with the H01 dataset analyses, Gillian Kay for scientific editing, the electron microscopy units (intra-departmental core facility at the Hebrew University Medical School and the Alexander Silberman Institute of Life Sciences) for technical support. This study was supported by the Leona M. and Harry B. Helmsley Charitable Trust (2015PG-ISL007); and the Israel Science Foundation (grants 1882/16 and 2402/16) to ABZ.

## Competing interests

**None**.

**Figure S1.**
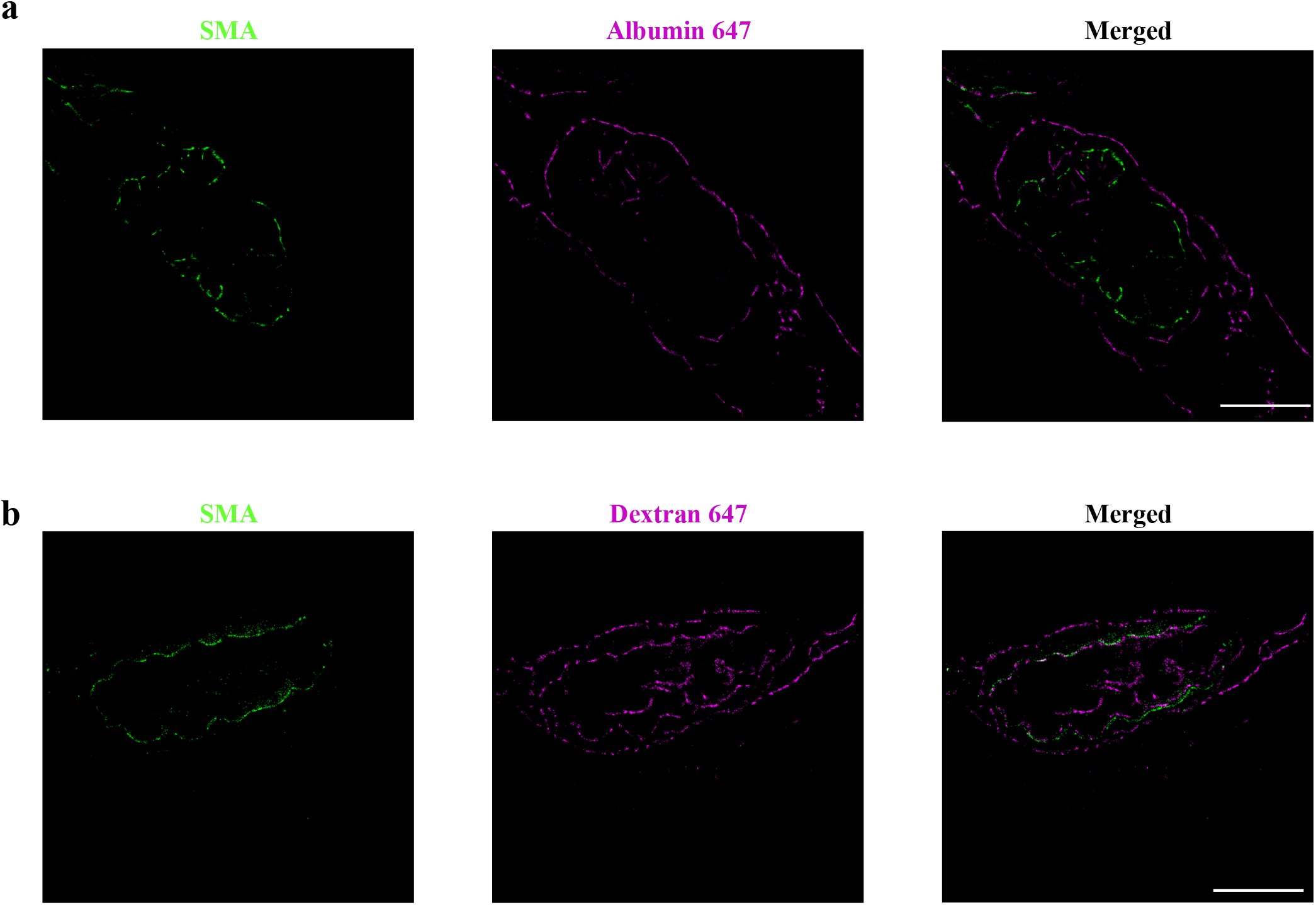
STORM imaging demonstrates super-resolution tracer permeability, validating unique barrier properties of the CNS arteriole-wall. Adult wild-type mouse cortical sections stained with SMA and imaged in STORM following tracer injections. Both albumin signals (**a)** and dextran signals (**b**) are found passed SMA markers. Scale bar 10 μm (n = 6 mice, 24 arterioles). Quantification is shown in Figure 2c.

**Figure S2.**
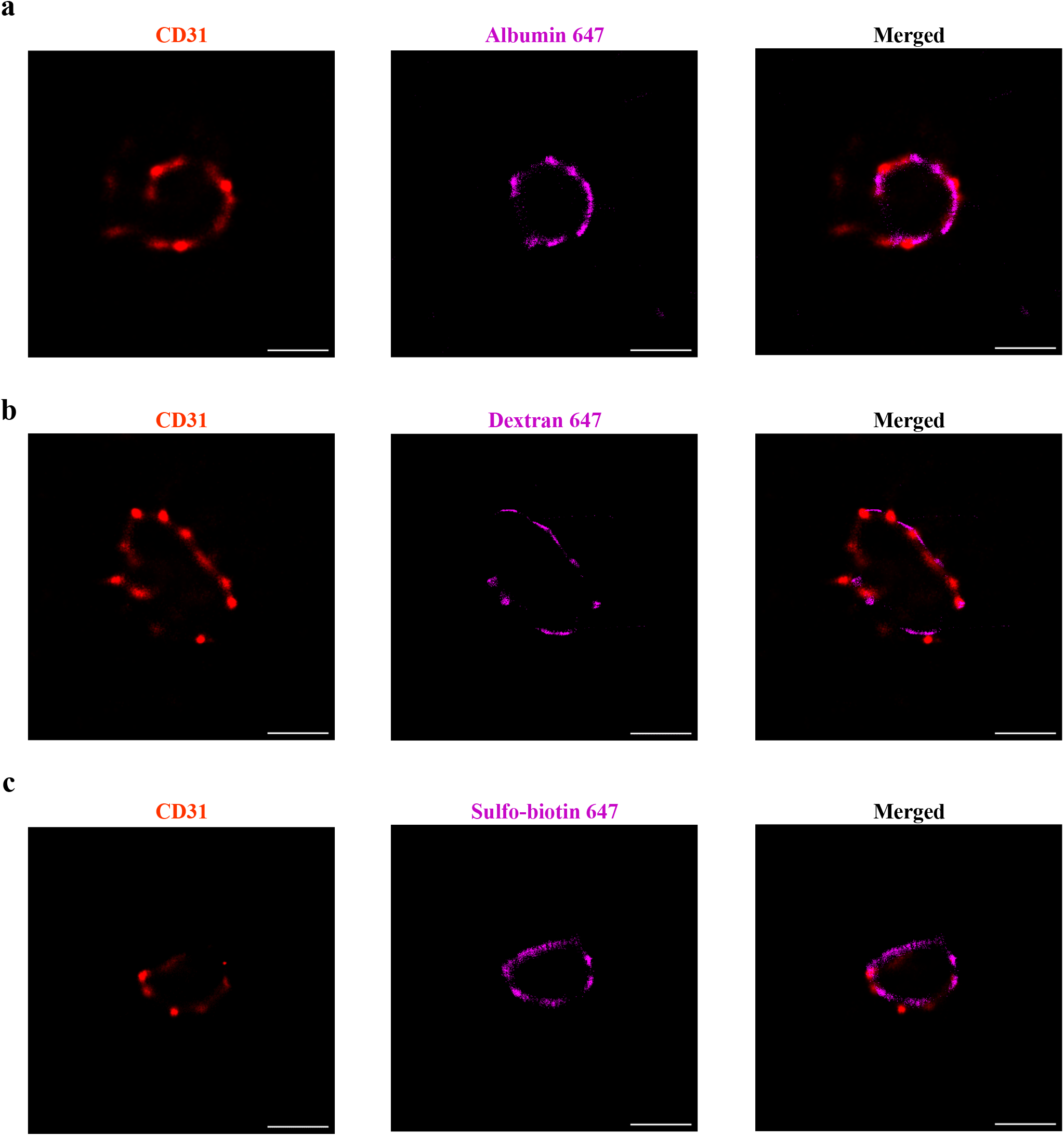
STORM imaging demonstrates super-resolution tracer permeability, confirming endothelial barrier properties of CNS capillaries. STORM imaging of wild-type adult mouse cortical capillaries, following tracer challenges. Sections were stained for an endothelial marker (CD31). Tracer challenge demonstrates that all three tracers are mostly co-localized with CD31 (**a**, albumin 647 (70 kDa), **b**, dextan 647 (10 kDa), and **c**, sulfo-biotin 647 (443 Da)). Scale Bar 10 μm (n = 9 mice, 27 capillaries).

